# Efficient parameterization of large-scale dynamic models based on relative measurements

**DOI:** 10.1101/579045

**Authors:** Leonard Schmiester, Yannik Schälte, Fabian Fröhlich, Jan Hasenauer, Daniel Weindl

## Abstract

**Motivation:** Mechanistic models of biochemical reaction networks facilitate the quantitative understanding of biological processes and the integration of heterogeneous datasets. However, some biological processes require the consideration of comprehensive reaction networks and therefore large-scale models. Parameter estimation for such models poses great challenges, in particular when the data are on a relative scale.

**Results:** Here, we propose a novel hierarchical approach combining (i) the efficient analytic evaluation of optimal scaling, offset, and error model parameters with (ii) the scalable evaluation of objective function gradients using adjoint sensitivity analysis. We evaluate the properties of the methods by parameterizing a pan-cancer ordinary differential equation model (*>* 1000 state variables, *>* 4000 parameters) using relative protein, phospho-protein and viability measurements. The hierarchical formulation improves optimizer performance considerably. Furthermore, we show that this approach allows estimating error model parameters with negligible computational overhead when no experimental estimates are available, providing an unbiased way to weight heterogeneous data. Overall, our hierarchical formulation is applicable to a wide range of models, and allows for the efficient parameterization of large-scale models based on heterogeneous relative measurements.

**Supplementary information:** Supplementary information are available at *bioRxiv* online. Supplementary code and data are available online at http://doi.org/10.5281/zenodo.2593839 and http://doi.org/10.5281/zenodo.2592186.

## 1 Introduction

In systems biology, mechanistic ordinary differential equation (ODE) models are widely used to deepen the understanding of biological processes. Applications range from the description of signaling pathways (Klipp *et al.*, 2005) to the prediction of drug responses (Hass *et al.*, 2017) and patient survival (Fey *et al.*, 2015). With the availability of scalable computational methods and increasing computing power, larger and larger models have been developed to capture the intricacies of biological regulatory networks more accurately (Bouhaddou *et al.*, 2018; Fröhlich *et al.*, 2018). In Fröhlich *et al.* (2018), we demonstrated how such a large-scale mechanistic model integrating various cancer-related signaling pathways is able to, e.g., predict the response of cancer cells to drug combinations based on measurements for single treatment responses, a task which is commonly not possible with statistical models. Overall, mechanistic models can pave the way to personalized medicine by integrating patient specific information, and thus creating virtual patients (Kühn and Lehrach, 2012; Ogilvie *et al.*, 2015).

Mechanistic ODE models usually contain parameters such as reaction rate constants and initial concentrations, which have to be inferred from experimental data. Parameter estimation for larger models is limited by (i) computational power for large numbers of required model simulations and gradient evaluations, as well as by (ii) the availability of data to infer parameter values. Scalable methods have been developed to address the problem of computational complexity, e.g. adjoint sensitivity analysis (Fujarewicz *et al.*, 2005; Lu *et al.*, 2012; Fröhlich *et al.*, 2017b) and parallelization (Penas *et al.*, 2015; Fröhlich *et al.*, 2018). Complementary, large-scale transcriptomics, proteomics and pharmacological datasets have been acquired and have been made publicly available in databases such as the Cancer Cell Line Encyclopedia (CCLE) (Barretina *et al.*, 2012), the Genomics of Drug Sensitivity in Cancer (GDSC) project (Eduati *et al.*, 2017) and the MD Anderson Cell Lines Project (MCLP) (Li *et al.*, 2017).

The available databases are rather comprehensive and cover already hundreds of cell-lines. Yet, those datasets are usually relative measurements and data often undergo some type of normalization, which has to be accounted for when linking mechanistic model simulations to the data. A commonly used approach is to introduce scaling and offset parameters in the model outputs (Weber *et al.*, 2011; Raue *et al.*, 2013; Degasperi *et al.*, 2017). However, this increases the dimensionality of the optimization problem and slows down optimization. Indeed, even a small number of scaling factors can result in a substantial drop of optimizer performance (Degasperi *et al.*, 2017). The precise reasons are yet to be understood.

To improve optimizer performance, Weber *et al.* (2011) developed a hierarchical optimization method which exploits the fact that for given dynamic parameters, the optimal scaling parameters can be computed analytically, which improved convergence and reduced computation time. The approach was generalized by Loos *et al.* (2018) to error model parameters and different noise distributions. However, the available approaches only considered scaling parameters, but not offset parameters. In addition, those approaches were not compatible with adjoint sensitivity analysis, but only with forward sensitivity analysis, which is computationally prohibitive for large-scale models.

Here, we (i) analyze the problems caused by the introduction of scaling factors and (ii) extend the hierarchical optimization method introduced by Loos *et al.* (2018) to be used in combination with adjoint sensitivity analysis. Furthermore, we derive the governing equations to not only include scaling parameters, but also offset parameters and the combination of both as well as error model parameters in the case of additive Gaussian noise. Our method is more general and achieves a better scaling behaviour than the existing ones (Weber *et al.*, 2011; Loos *et al.*, 2018). We apply it to estimate parameters for the large-scale pan-cancer signaling model from Fröhlich *et al.* (2018). First, we use simulated relative and absolute data to compare the performance of the standard and the novel hierarchical approach and to demonstrate the loss of information associated with using only relative data. Second, we use measured data to estimate model parameters, compare the performance of different optimization algorithms, and show how the performance of each of them improves with our hierarchical optimization approach.

## 2 Materials and Methods

### 2.1 Mechanistic modeling

We consider ODE models of biochemical processes of the form

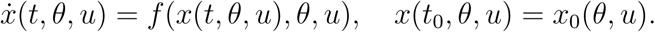

The state vector 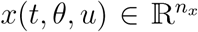 denotes the concentrations of involved species, the vector field 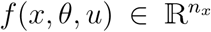 describes the temporal evolution of the states, the vector 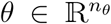 unknown parameters, the vector 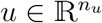 differential experimental conditions, and 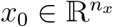 the parameter-and condition-dependent states at initial time *t*_0_.

An observation function *h* maps the system states to observables 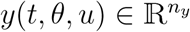, via

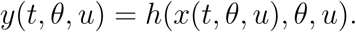

Experimental data 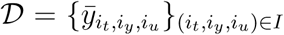 corresponding to the observables are time-discrete and subject to measurement noise 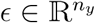,

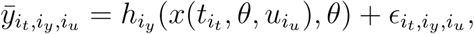

indexed over a finite index set *I* of time points *i*_*t*_, observables *i*_*y*_, and experimental conditions *i*_*u*_. We assume the measurement noise to be normally distributed and independent for all datapoints, 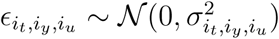.

### 2.2 Relative measurements

Frequently, experiments provide measurement data only in a relative form, in arbitrary units, rather than as absolute concentrations. Thus, to compare model and data, the observables need to be rescaled. While the rescaling is usually incorporated in *h* and *θ*, here we use an explicit formulation. Since these cover a broad range of measurement types, we assume that we have scaling factors *s* and offsets *b* such that simulations and measured data are related via

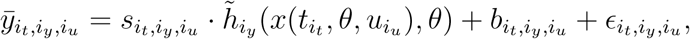

in which 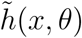 denotes the mapping to unscaled observables.

Scaling factors 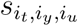 and offsets 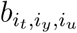, but also noise parameters 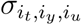, in the setting considered here the standard deviations of Gaussian distributions, are often shared between some datapoints, e.g. for time series measurements, or for data taken under the same experimental conditions. In the following, we summarize all different scaling, offset and noise parameters in vectors 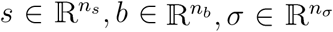, respectively, and refer to them as static parameters, to distinguish them from the original parameters *θ*, henceforth called dynamic parameters, since they affect the dynamics of the simulated states. The static parameters are often unknown and thus have to be estimated along with the dynamic parameters.

### 2.3 Parameter estimation problem with relative data

To infer the unknown parameters *θ, s, b*, and *σ*, we maximize the likelihood

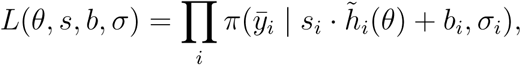

of observing the experimental data 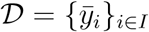 given parameters *θ, s, b, σ*, where for simplicity of presentation we employ a general index set *i ∈ I* over time points, observables, and experimental conditions. *π* denotes the conditional probability of observing 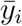 given simulation 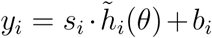 and noise parameters *σ*_*i*_. For Gaussian noise, we have

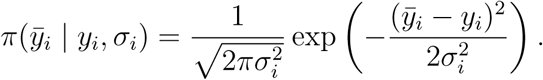

Instead of maximizing *L* directly, it is equivalent and numerically often preferable to minimize the negative log-likelihood min_*θ,s,b,σ*_ *J* (*θ, s, b, σ*) with *J* = −log *L*. Assuming Gaussian noise, *J* becomes

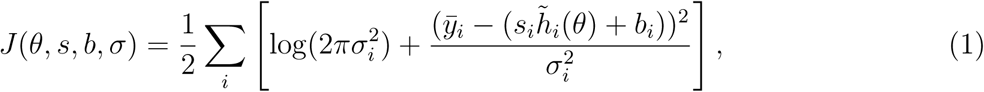

which will henceforth be referred to as objective function.

### 2.4 Hierarchical optimization

In this section, we generalize the hierarchical optimization approach introduced by Loos *et al.* (2018) to (1), allowing for scaling, offset, and noise parameters simultaneously, and we outline how hierarchical optimization can be combined with adjoint sensitivities.

The standard approach to handle the static parameters is to consider the extended parameter vector (*θ, s, b, σ*) and to optimize all its elements simultaneously. However, the increased dimension makes the optimization problem in general harder to solve. Instead, we can make use of the specific problem structure of (1) more efficiently by splitting the optimization problem into an outer problem where we optimize the dynamic parameters *θ*, and an inner problem where we optimize the static parameters *s, b*, and *σ*, conditioned on *θ*. That is, we compute

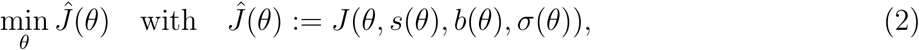

in which

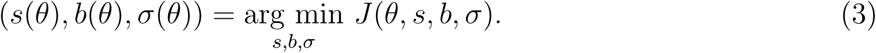

It can be shown that global optima of the standard optimization problem are preserved in the hierarchical problem.

#### 2.4.1 Analytic expressions for the optimal scaling, offset, and noise parameters

In general, an inner optimization problem like (3) needs to be solved numerically. However, under certain conditions one can give analytic expressions for the optimal static parameters, which renders solving the inner problem computationally very cheap. The analytic expressions are based on evaluating the necessary condition for a local minimum in *s, b, σ* given *θ*,

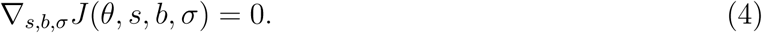

Here, we extend the available results by Weber *et al.* (2011) and Loos *et al.* (2018).

We define index sets 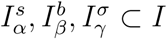 for *α* = 1, *…*, *n*_*s*_, *β* = 1, *…*, *n*_*b*_, *γ* = 1, *…*, *n*_*σ*_, with *n*_*s*_, *n*_*b*_ and *n*_*σ*_ indicating the number of scaling, offset and noise parameters. The index sets indicate which datapoints share static parameters, e.g., all datapoints 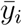 with 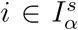 share a scaling parameter. In order to derive analytic formulas, we will in the following assume that 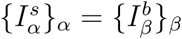, i.e., that scaling and offset parameters are shared among the same datapoints, and that for all *α* there exists *γ* such that 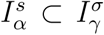, i.e., that datapoints sharing the scaling (and offset) parameter share also the noise parameter. The results are also flexible enough to allow any of *s, b*, and *σ* to be fixed or estimated as dynamic parameters, in which case the above assumptions can be alleviated. For an extended discussion and derivations of the below formulas see the Supplementary Information, Section 3.

First, we consider single scaling parameters *s*_*α*_ and offset parameters *b*_*β*_. Without loss of generality, we reduce the objective (1) to only include relevant summands. Then, (4) yields

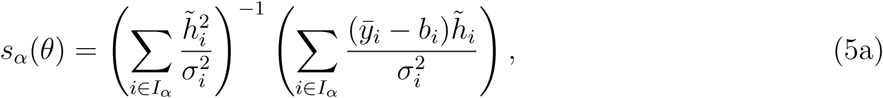

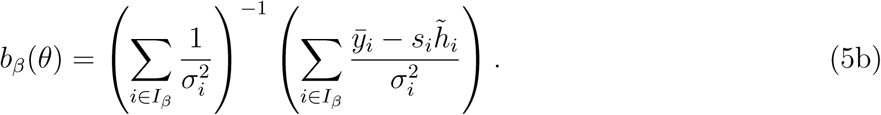

If either the *s*_*i*_ or *b*_*i*_ are no static parameters, we are done by just inserting those values in the respective other formula. If both are to be optimized as static parameters, in which case by assumption *s*_*i*_ *≡ s*_*α*_, *b*_*i*_ *≡ b*_*β*_, we can proceed by inserting (5a) into (5b), which yields non-interdependent formulas, see the Supplementary Information, Section 3.1. Note that the noise parameters drop out of the formulas if all values coincide, as is our assumption in the case that we want to estimate the noise parameters hierarchically as well. Thus, in either case *s*_*α*_(*θ*) and *b*_*β*_(*θ*) can now be readily computed. Note that for the special case *b* = 0 we recover the formula from Loos *et al.* (2018).

Second, for a given single noise parameter *σ*_*γ*_, we consider without loss of generality an objective function (1) reduced to indices *I*_*γ*_, while *s*_*i*_ and *b*_*i*_ can be arbitrary. The objective considered here will typically contain multiple sums of the type discussed for the scalings and offsets. As *s* and *b* are known already at this stage, (4) immediately gives

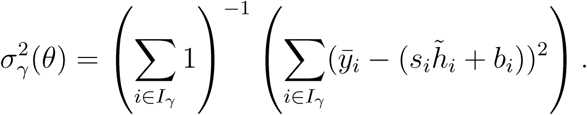

Note that a problem occurs when the rescaled simulations match the measured data exactly, since then *σ*^2^ = 0. In this case, the noise parameter and thus the objective function is unbounded in the standard and the hierarchical formulation, so that measures to deal with this case have to be taken, e.g. by specifying a lower bound for *σ*_*γ*_.

Inspection of the Hessian 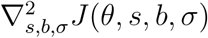 shows that the found stationary points indeed are minima (see Supplementary Information, Section 3).

#### 2.4.2 Combining hierarchical optimization and adjoint sensitivity analysis

In optimization, the objective function gradient is of considerable value, because it gives the direction of steepest descent in the objective function landscape. Recent studies indicated that optimization methods using gradients tend to outperform those which do not (Schälte *et al.*, 2018; Villaverde *et al.*, 2018). In Loos *et al.* (2018), hierarchical optimization was performed using objective gradients computed via forward sensitivity analysis. However, for large-scale models adjoint sensitivity analysis has shown to be orders of magnitude faster (Fröhlich *et al.*, 2018), because essentially here the evaluation of state sensitivities is circumvented by defining an adjoint state 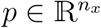 which does not scale in the number of parameters (Fröhlich *et al.*, 2017b).

Whether hierarchical optimization can be combined with adjoint sensitivity analysis so far remained unclear. The problem with applying adjoint sensitivity analysis is that, unlike the forward sensitivity equations, the adjoint state depends on the data and the scaled observables and thus requires knowledge of the static parameters. Therefore, the approaches by Weber *et al.* (2011) and Loos *et al.* (2018) of first simulating the state trajectory *x*(*t, θ, u*) as well as all required sensitivities, and then computing optimal static parameters in order to compute *Ĵ* and *∇Ĵ* without further simulations, are not applicable.

To combine hierarchical optimization and adjoint sensitivity analysis, we derived the scheme illustrated in Figure 1. Conceptually, we postpone the evaluation of the adjoint state to after the computation of the optimal static parameters. As the derivatives of the objective function with respect to the optimal static parameters are zero, i.e. *∇*_*s,b,σ*_*J* = 0, since we solve the inner sub-problem exactly, we can prove that this scheme provides the correct objective function gradient *∇Ĵ*. For a more detailed discussion and derivation of the adjoint-hierarchical approach, we refer to the Supplementary Information, Section 2. An overview over the properties of the different hierarchical optimization approaches is provided in the Supplementary Information, Table S1.

**Figure 1:**
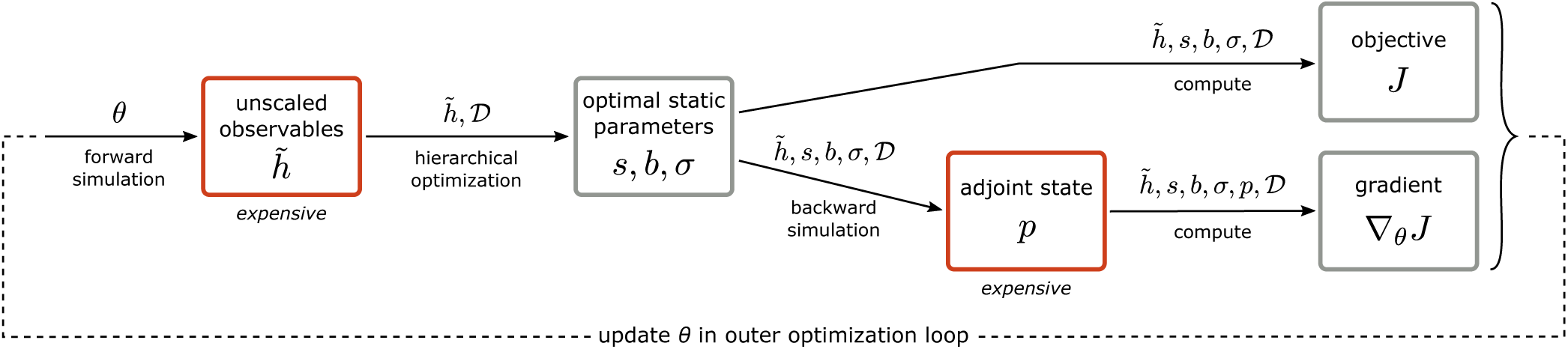
Illustration of the hierarchical optimization scheme using adjoint sensitivities. In the outer loop, *θ* is updated by some iterative optimization scheme. Here, the solution of the inner problem is shown in detail. The red boxes involve the simulation of ODEs and are thus usually computationally more expensive. If the gradient is not required in some optimizer iteration, the adjoint and gradient steps can be omitted.

### 2.5 Implementation

We implemented the proposed method in MATLAB and C++. A custom parallelized objective function implementation was used to decrease the wall time (see Supplementary Information, Sections 4.5.2 and 4.5.3). The modular implementation can easily be adopted to work with other Systems Biology Markup Language (SBML, Hucka *et al.* (2003)) models. Model simulation and gradient evaluation using the proposed scheme were performed using AMICI (Fröhlich *et al.*, 2017a). Parameter estimation was performed using multi-start local optimization. The starting points were sampled from a uniform distribution. The initial dynamic parameters were identical for the standard and hierarchical optimization, where initial static parameters only had to be chosen for the standard approach. We considered different local optimization methods (see Section *Results*) and ran all for a maximum of 150 iterations (see Supplementary Information, Section 4.5.1 for more details). The complete code and data are available at http://doi.org/10.5281/zenodo.2593839 and http://doi.org/10.5281/zenodo.2592186.

## 3 Results

In this study, we considered the pan-cancer signaling pathway model developed by Fröhlich *et al.* (2018). This model comprises 1396 biochemical species (1228 dynamic states and 168 constant species) and 4232 unknown parameters, and can be individualized to specific cancer cell-lines using genetic profiles and gene expression data. Fröhlich *et al.* (2018) demonstrated a promising performance of the model in drug response prediction, but molecular insights were limited by non-identifiabilities. Motivated by these results, we set out to parameterize this model using additional data.

### 3.1 Mapping multiple datasets to a large-scale model of cancer signaling

For model calibration, we considered two datasets. *Dataset 1* is the training data studied by Fröhlich *et al.* (2018). These are viability measurements for 96 cancer cell-lines in response to 7 drugs at 8 drug concentrations available in the CCLE (Barretina *et al.*, 2012). The viability measurements are normalized to the respective control. To account for this normalization, Fröhlich *et al.* (2018) simulated the model for the treated condition and the control, and the simulations were then divided by each other. This corresponds to the method proposed by Degasperi *et al.* (2017). However, this approach is not applicable if multiple observables need to be considered, e.g. when incorporating additional data types, or when more complex data normalizations are applied. Therefore, we reformulated the model output and replaced the normalization with the control by a cell-line specific scaling 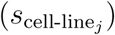. This yields the observation model

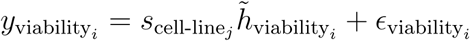

with *i* indexing the datapoints belonging to cell-line *j*. The measurement noise is assumed to be normally distributed, 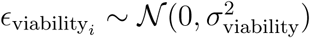.

We complemented the viability measurements employed with molecular measurements to refine the parameter estimates. *Dataset 2* contains reverse phase protein array (phospho-)proteomic data for various cancer cell-lines taken from the MCLP (Li *et al.*, 2017). We developed a pipeline which (i) maps the measured protein levels to the state variables of the model and (ii) employs the mapping to construct observables (see Supplementary Information, section 4.1 for more details). We identified 32 proteins and 16 phosphoproteins measured that were also covered by the model. In total, 54 out of the 96 considered cell-lines were included in the MCLP (*dataset 2* in Table 1). In the MCLP database, measurements are normalized across cell-lines and across all proteins by subtracting the respective median from the log2-transformed measured values (see *Level 4 data* in https://tcpaportal.org/mclp/#/faq). Therefore, we included one cell-line specific offset 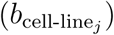 and one protein specific offset 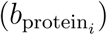, yielding the observation model

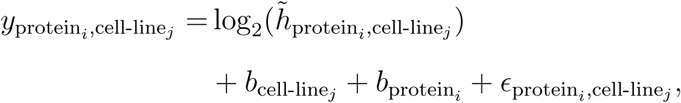

normally distributed measurement noise 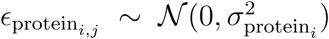 and the simulated absolute protein level

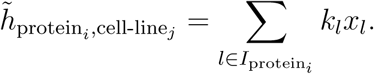

**Table 1:** Datasets used for parameter estimation. The number of static parameters of certain classes is indicated, followed by the number of parameters which are computed analytically in the hierarchical setting in parentheses.

|  | Dataset 1 (CCLE) | Dataset 2 (MCLP) |
| --- | --- | --- |
| # datapoints | 5281 | 1799 |
| # cell-lines | 96 | 54 |
| # observables | 1 | 48 |
| # scalings | 96(96) | 0 |
| # offsets | 0 | 102(48) |
| # noise parameters | 1(1)* | 48(48) |

\*The noise parameter is set to one if dataset 1 is considered individually.

The index set 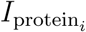 refers to the species that include protein_*i*_ and *k*_*l*_ is the respective stoichiometric multiplicity.

The integration of viability and molecular measurements provides information on two different levels, which potentially improves the reliability of the model. However, it requires a substantial number of observation parameters (Table 1).

### 3.2 Evaluation of standard and hierarchical optimization using simulated data

A priori it is not clear which influence scaling, offset and noise parameter have on optimizer performance. However, Degasperi *et al.* (2017) observed in two examples that the use of scalings lead to inferior optimizer behaviour compared to the normalization-based approach which was also used by Fröhlich *et al.* (2018). Thus, before estimating parameters using real measured data from CCLE and MCLP, we first used simulated data. To get realistic data, we simulated the model for the same experimental conditions that were provided in *dataset 1* and added normally distributed noise to the simulations (see Supplementary Information, Section 4.7). The simulation of experimental data allowed us to (i) compare the goodness-of-fit of estimated and true parameter and to (ii) assess the information associated with relative data.

#### 3.2.1 Hierarchical optimization facilitates convergence

To compare standard and hierarchical optimization, we employed both approaches for the analysis of simulated, noisy relative data. For local optimization we employed the Interior Point OPTimizer (Ipopt) (Wächter and Biegler, 2006; HSL, 2019). As metric we considered the Pearson correlations between data and simulation for each of the optimized parameter vectors and the true parameter vector. The Pearson correlation reflected the objective function value (Supplementary Information, Figure S1) but was easier to interpret.

The hierarchical optimization achieved substantially better correlations between simulation and data than the standard optimization (Figure 2A). Furthermore, variability between different local optimization runs was reduced. Indeed, all but two optimizer runs using hierarchical optimization achieved correlations similar to the correlation observed for the true parameters, indicating a good model fit and – in contrast to the standard optimization – a good convergence. No run reached significantly better values than the true parameter vectors which would have indicated over-fitting.

**Figure 2:**
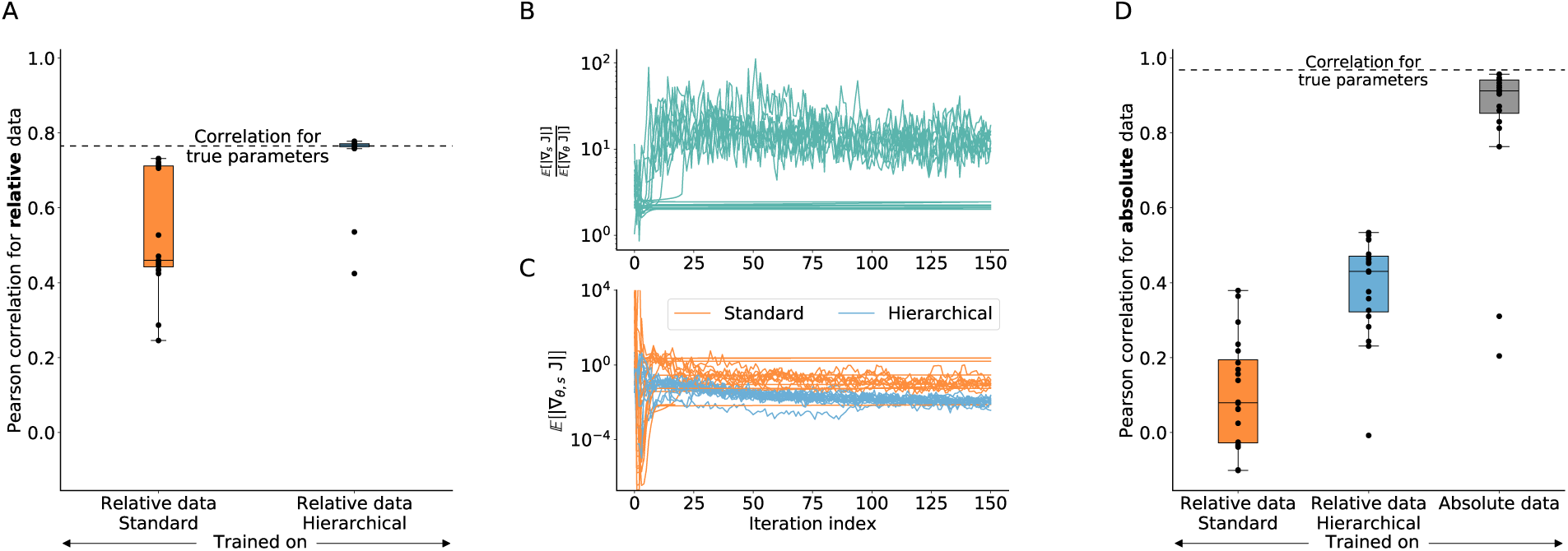
Convergence of standard and hierarchical optimization. Parameter estimation results using a simulated version of *dataset 1* from Table 1 with the Ipopt optimizer. For all evaluations 20 optimizer runs were performed. **A**: Pearson correlation of relative training data and corresponding model simulation after training on relative data using standard and hierarchical optimization. Dashed line shows the correlation that is achieved using the true parameters used to generate the training data. **B**: Ratio of the average gradient contribution for scaling parameters against dynamic parameters using the standard optimization for all optimizer runs along their trajectory. **C**: Expected gradient for standard and hierarchical optimization. Only the parameters, that were optimized numerically, were taken into account. **D**: Pearson correlation of absolute data and corresponding model simulation after training on (left & middle) relative data and (right) on absolute data.

#### 3.2.2 Scalings have a pronounced influence on the objective function value

Hierarchical optimization decreases the effective dimension of the optimization problem. However, as the number of parameters decreases for the considered problem only by 2% – this does not explain the substantially improved convergence – the scaling factors might be particularly relevant. To assess this, we evaluated the average absolute values of the objective function gradient for scaling parameters (*E*[*|∇*_*s*_*J|*]) and dynamic parameters (*E*[*|∇*_*θ*_*J|*]). Indeed, the evaluation of the ratio (*E*[*|∇*_*s*_*J|*]*/E*[*|∇*_*θ*_*J |*]) revealed that the objective function is usually 10 times more sensitive with respect to scaling parameters than dynamic parameters (Figure 2B). This indicates the presence of two separate timescales in the continuous representation of the optimization problem (Wibisono *et al.*, 2016), which suggests that the optimization problem is stiff. As standard optimization methods correspond to explicit solving schemes of the continuous optimization problem, the stiffness could explain the problem encountered for the standard approach. Vice versa, it explains the improvement achieved using hierarchical optimization, where the gradient contribution of the scalings is zero. Accordingly, the average gradient for the hierarchical optimization is small compared to the standard optimization (Figure 2C).

An inspection of the optimizer trajectories revealed that for the standard optimization some optimizer runs show flat trajectories of the objective function, while still having a comparably large gradient (Figure 2C and Supplementary Information, Figure S2). For these runs, the contribution of the scalings became small (flat lines in Figure 2B), which might be due to a valley in the objective function landscape defined by the scaling parameters, where the optimizer got stuck. Such valleys are eliminated in the hierarchical optimization.

#### 3.2.3 Normalization results in information loss

To assess the influence of information loss associated with the use of relative data, we performed optimization using simulated absolute data. For comparison, we predicted the absolute values using the parameters inferred with relative data (see Supplementary Information, Section 4.7.3). As expected, we found that the prediction of absolute data from relative data yields a correlation far from one (Figure 2D), implying that information is lost in the normalization process. Interestingly, hierarchical optimization again outperformed standard optimization. A potential reason is that the improved convergence of the optimizer allows for the extraction of more information from the relative data.

### 3.3 All tested local optimization methods profit from hierarchical formulation

To provide a thorough comparison of the performance of standard and hierarchical optimization, we assessed it for different local optimization algorithms on the measured viability data (*dataset 1*). We considered four commonly used or open-source optimizers: Ipopt (Wächter and Biegler, 2006; HSL, 2019), Ceres (Agarwal *et al.*, 2019), sumsl (Gay, 1983), and fmincon (Mathworks, 2019). These optimizers use different updating schemes, e.g. based on line-search or trust-region methods.

We assessed the performance by studying the evolution of objective function values over computation time and optimizer iterations. Given the same computational budget, the hierarchical optimization consistently achieved better objective function values for all considered optimization algorithms and for almost all runs (Figure 3A and Supplementary Information, Figure S3). Furthermore, the objective function at the maximum number of iterations was substantially better for hierarchical optimization than standard optimization, and there was in general a lower variability (Figure 3B). Given this result, we determined the computation time required by the hierarchical optimization to achieve the final objective function value of the standard optimization and computed the resulting speed-up (Figure 3C). Except for one start of Ipopt, the hierarchical optimization was always faster with a median speed-up between one and two orders of magnitude. Since a single local optimization run required several thousand hours of computation time, the efficiency improvement achieved using hierarchical optimization is crucial.

**Figure 3:**
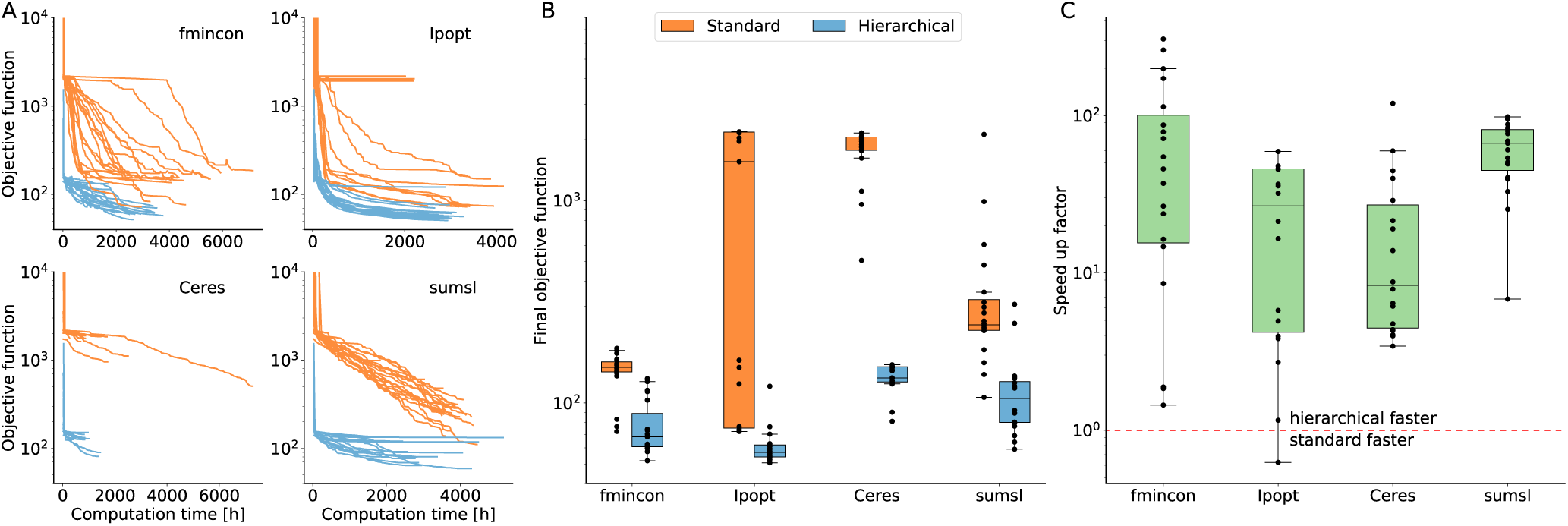
Computational efficiency of standard and hierarchical optimization for multiple optimization algorithms. **A**: Optimizer trajectories for fmincon, Ipopt, Ceres and sumsl using standard and the hierarchical optimization. Since the noise parameter was set to 1 for these runs, the constant term in the objective function was omitted. *dataset 1* from Table 1 was used. Fmincon runs were performed on different systems and using a different implementation than the other optimizers, so that absolute computation times are not comparable. **B**: Boxplots of final objective function values obtained after 150 iterations by the different optimizers using standard and hierarchical optimization. **C**: Speed-up of the hierarchical optimization compared to the standard optimization. The speed-up is defined by the computation time the hierarchical optimization needs to find the final objective function value of the standard optimization for every local optimization (or vice versa if the standard optimization finds a better final value). The dashed red line shows the point, where standard and hierarchical are equally fast.

As the performance of optimization algorithms has so far mostly been evaluated for ODE models with tens and hundreds of unknown parameters (Villaverde *et al.*, 2018; Hass *et al.*, 2019), we used our results for a first comparison on a large-scale ODE model. We found that for the considered problem (i) Ceres always stopped prematurely, (ii) sumsl progressed (at least for the standard optimization) slower than Ipopt and fmincon, and (iii) fmincon and Ipopt reached the best objective function values and appeared to be most efficient (Figure 3A, B).

### 3.4 Hierarchical optimization enables integration of heterogeneous data

As the information about molecular mechanisms provided by viability measurements (*dataset 1*) are limited, we complemented it using the (phospho-)protein measurements (*dataset 2*). An unbiased weighting was ensured by introducing error model parameters (i.e., standard deviations) for the individual observables and estimating them along with the remaining parameters. In hierarchical optimization, (i) the error model parameters, (ii) the cell-line specific scaling of the viability measurements and (iii) the observable-specific offsets of the log-transformed protein measurements are optimized analytically (Table 1). The analytic optimization of the cell-line specific offsets of the log-transformed protein measurements is not supported by the approach as the error model parameters and the offsets have to share the same datapoints.

We performed multi-start local optimization for the combined dataset using Ipopt. Again, the hierarchical optimization was computationally much more efficient and reached better objective function values than the standard optimization (Figure 4A). For the standard optimization, all starts yielded objective function values of approximately 10^4^. For the hierarchical optimization, we observed runs yielding objective values similar to those for standard optimization denoted by Group 1, with *J ≈* 10^4^ as well as runs which provided much better objective function values, i.e. Group 2, *J <* 3 *×* 10^3^. The optimized parameter vectors obtained using standard optimization runs and hierarchical optimization runs in Group 1 were able to fit the viability measurements but failed to describe the protein data (Figure 4B). In contrast, the optimized parameter vectors obtained using hierarchical optimization runs in Group 2 show a good fit for viability and most protein measurements (Figure 4B). Accordingly, only hierarchical optimization runs managed to balance the fit of the datasets, thereby achieving an integration and a better overall description of the data.

**Figure 4:**
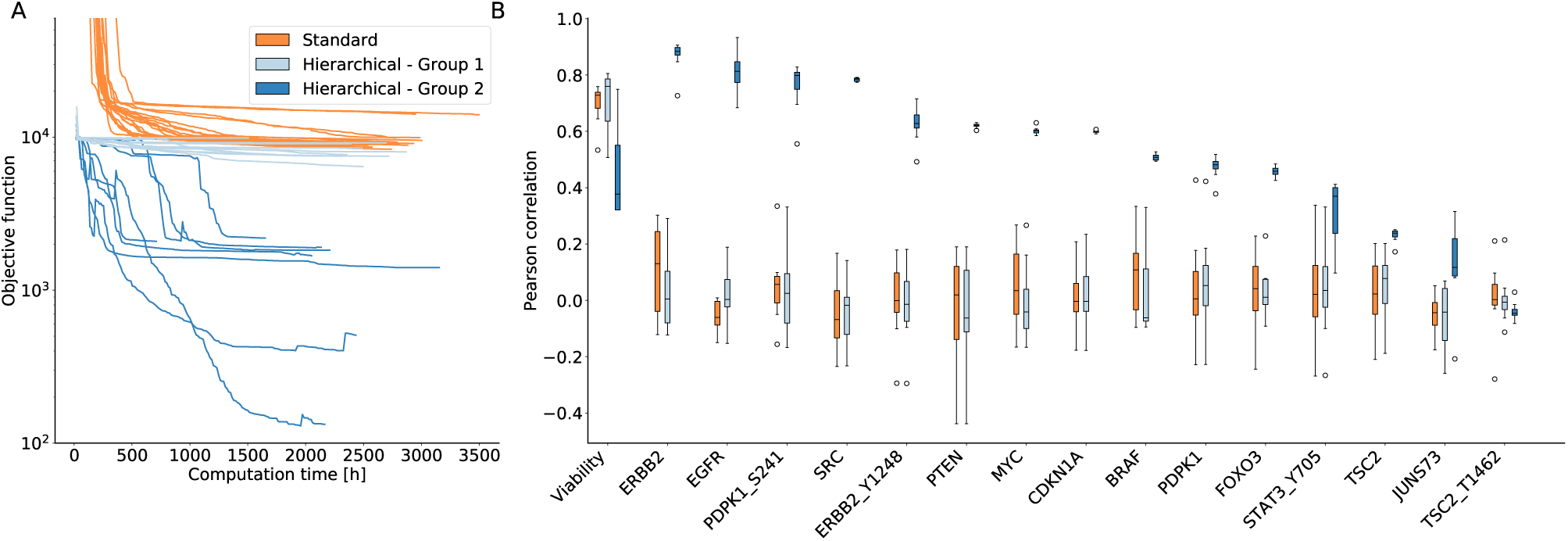
Integration of heterogeneous data using hierarchical optimization. **A**: Optimizer trajectories for standard and hierarchical optimization with *dataset 1 and 2* from Table 1 using Ipopt. The two Groups found by the hierarchical optimization are indicated by different shades of blue. **B**: Pearson correlations for all observables with at least 55 datapoints for all runs of the standard optimization and for the two groups found by the hierarchical optimization. For all observables, see Supplementary Information, Figure S4.

In summary, the adjoint-hierarchical approach outperformed in all regards standard optimization. Compared to forward-hierarchical approaches (Weber *et al.*, 2011; Loos *et al.*, 2018) a speedup of roughly three orders of magnitude is achieved. The computation time of one gradient evaluation with forward sensitivities is in the same order as one full optimization, with the here used settings, applying adjoints (see Supplementary Information, Section 4.3 for an estimate of the computation time).

## 4 Discussion

Parameterization of large-scale mechanistic models is a challenging task requiring new approaches. Here, we combine the concept of hierarchical optimization (Weber *et al.*, 2011; Loos *et al.*, 2018) with adjoint sensitivities (Fujarewicz *et al.*, 2005; Lu *et al.*, 2012; Fröhlich *et al.*, 2017b). This is crucial when parameterizing large-scale models for which the use of forward sensitivities is computationally prohibitive. Additionally, we derived more general formulas for hierarchically optimizing a combination of scaling and offset parameters as well as noise parameters.

We demonstrated the advantages of hierarchical optimization using a recently published large-scale pan-cancer model. We obtained median speed-ups of more than one order of magnitude as compared to the conventional approach, irrespective of the employed optimizer. Given that the overall computation time is thousands of CPU hours, this improvement is substantial. While previous studies had already shown a reduced convergence rate when calibrating models to relative data (Degasperi *et al.*, 2017), we identified the large gradients with respect to the scalings as a possible explanation and established a flexible and easy way to circumvent them. The numerical stiffness which can arise from this for numerical optimization methods is the first conceptual explanation of the large improvements achieved by hierarchical methods (Weber *et al.*, 2011; Loos *et al.*, 2018).

In addition to the methodological contribution, we provide here the first proof-of-principle for the integration of multiple datasets using large-scale mechanistic models of cancer signaling. We showed for the example of viability and (phospho-)proteomic measurements that our optimization approach facilitates (i) data integration – where other methods failed – and (ii) an easy weighting of datasets. This is possible without computational overhead. The optimized noise parameters provide estimates for the measurement noise when no or only low numbers of replicates are available, as it is the case in many large-scale databases (e.g. CCLE and MCLP).

In this study, we used hierarchical optimization to estimate individual static parameters per observable. However, measurements may require multiple scaling and offset parameters per observable (e.g the protein observables considered here), as well as arbitrary combinations thereof. The current hierarchical framework cannot account for such settings. An extension to efficiently estimate all such parameters would thus presumably yield an even improved performance. Similarly, extending the optimization approach to other noise models would be of interest, even when the inner subproblem lacks an analytical solution. Of particular interest are distributions that are more robust to outliers, while still maintaining the good optimization convergence (Maier *et al.*, 2017).

Large-scale mechanistic models are of high value for systems biomedicine, since, as opposed to machine learning methods, they allow for mechanistic interpretation, analysis of latent variables and extrapolation to unseen conditions (Baker *et al.*, 2018; Fr ö hlich *et al.*, 2018). While this study is a proof-of-concept for the integration of heterogeneous datasets, for future biology-driven analyses it will be valuable to include additional molecular measurements to improve the predictive power and the mechanistic interpretation of the model. With the advance of high-throughput technologies, more and more such large-scale datasets have been published. For example, the cancer proteomic atlas (TCPA) (Li *et al.*, 2013) or the datasets provided by Frejno *et al.* (2017) or Gholami *et al.* (2013) constitute rich sources of training data for future analyses. Our hierarchical optimization now allows for a much more efficient calibration of large-scale mechanistic models using heterogeneous datasets.

## Acknowledgements

The authors declare no conflict of interest.

## Funding

This work was supported by the German Research Foundation (Grant No. HA7376/1-1; Y.S.), the German Federal Ministry of Education and Research (SYS-Stomach; Grant No. 01ZX1310B; J.H.), and the European Union’s Horizon 2020 research and innovation program (CanPathPro; Grant No. 686282; F.F., J.H., D.W.). Computer resources for this project have been provided by the Gauss Centre for Supercomputing / Leibniz Supercomputing Centre under grant pr62li.

## Author Contributions

Y.S., J.H. derived the theoretical foundation; D.W., F.F, L.S., Y.S., wrote the implementations; D.W., L.S. performed the case study. All authors discussed the results and conclusions and jointly wrote and approved the final manuscript.

