## Supplementary Information for "Efficient parameterization of large-scale dynamic models based on relative measurements"

### 1 Overview

In the main manuscript, we discuss the analytic hierarchical optimization of scaling, offset, and noise parameters in a particular setting assuming additive Gaussian measurement noise. Here, we provide some background information and a more general description (Section 2). For the specific situation of objective functions based on ordinary differential equation (ODE) models, we outline for time-discrete measurements the adjoint approach of computing the objective gradient (Section 2.3). We describe how hierarchical optimization can be combined with adjoint sensitivities for efficient gradient-based optimization of large-scale problems (Section 2.4). Then, we discuss in detail the derivation of analytic expressions for scaling, offset, and noise parameters assuming an additive Gaussian noise model (Section 3). Last, we give details on the case study and its implementation (Section 4).

### 2 Hierarchical gradient-based optimization of ODE models

A common situation in systems biology is an ODE model of the form (this is the same as in the main manuscript, included here just for completeness of discussion)

$$\begin{aligned} \dot{x}(t,\theta,u) &= f(x(t,\theta,u),\theta,u), \qquad x(t_0,\theta,u) = x_0(\theta,u), \\ y(t,\theta,u) &= h(x(t,\theta,u),\theta,u). \end{aligned} \tag{1}$$

Here, the state vector $x(t,\theta,u) \in \mathbb{R}^{n_x}$ describes the involved species concentrations. The vector field $f : \mathbb{R}^{n_x} \times \mathbb{R}^{n_\theta} \times \mathbb{R}^{n_u} \to \mathbb{R}^{n_x}$ describes the temporal evolution of the states. The vector $x_0(\theta,u)$ denotes the initial states at time $t_0$. The vector $\theta \in \mathbb{R}^{n_\theta}$ denotes unknown parameters like reaction rates or initial concentrations, whereas $u \in \mathbb{R}^{n_u}$ encodes (possibly time-dependent) differential experimental conditions like drug doses, which are given as an input to the system, and need not be inferred. We first discuss the general problem, and link to hierarchical optimization later. The function $h : \mathbb{R}^{n_x} \times \mathbb{R}^{n_\theta} \times \mathbb{R}^{n_u} \to \mathbb{R}^{n_y}$ maps the system state to observables $y(t,\theta,u) \in \mathbb{R}^{n_y}$, where typically $n_y < n_x$ since not all states can be observed. These observables correspond to measured experimental data

$$\mathcal{D} = \{\bar{y}_{i_t,i_y,i_u}\}_{(i_t,i_y,i_u)\in I}$$

for some index set $I$ over time points $1 \le i_t \le n_t$, observables $1 \le i_y \le n_y$, and experiment-specific conditions $1 \le i_u \le n_u$ (where we w.l.o.g. include possible experimental or technical replicates in the experimental conditions). When the origin of a specific data point does not matter, we will simplify the notation to an arbitrary index $i \in I$. The measurements are linked to the states via

$$\bar{y}_{i_t,i_y,i_u} = h_{i_y}(x(t_{i_t},\theta,u_{i_u}),\theta,u_{i_u}) + \epsilon_{i_t,i_y,i_u}$$

for some noise model $\varepsilon_{i_t,i_y,i_u}$, a random variable. This noise model can again be subject to parameters, denoted $\sigma = \sigma(\theta)$ here, possibly parameter-dependent. Here, we will consider Gaussian noise $\varepsilon_{i_t,i_y,i_u} \sim \mathcal{N}(0,\sigma^2_{i_t,i_y,i_u})$, the most commonly used noise model. We assume the noise on all data points to be independent, however different noise models, e.g. for different conditions, are possible. To infer the unknown parameters $\theta$, we maximize the likelihood

$$L(\theta) = \prod_{i_t,i_y,i_u} \pi(\bar{y}_{i_t,i_y,i_u} \mid h_{i_y}(t_{i_t},\theta,u_{i_u}),\sigma)$$

of observing the measured data $\mathcal{D}$ given parameters $\theta$ under the noise model $\pi$. To increase convexity and for numeric reasons, usually we minimize the negative log-likelihood $J(\theta) := -\log L(\theta)$ instead. It takes the general form

$$J(\theta) = -\sum_{i_t,i_y,i_u} \log \pi(\bar{y}_{i_t,i_y,i_u} \mid y_{i_y}(t_{i_t},\theta,u_{i_u}),\sigma).$$

In particular, assuming Gaussian measurement noise, we get

$$J(\theta) = \sum_{i_t,i_y,i_u} \left[ \log(2\pi\sigma^2_{i_t,i_y,i_u}) + \frac{(\bar{y}_{i_t,i_y,i_u} - y_{i_y}(t_{i_t},\theta,u_{i_u}))^2}{\sigma^2_{i_t,i_y,i_u}} \right]. \tag{2}$$

#### 2.1 Gradient-based optimization

When we want to optimize $J$ w.r.t. the parameters $\theta$, we usually employ an iterative gradient-based scheme, assuming our objective is sufficiently smooth. That is because the gradient $\nabla_\theta J$ gives the direction of steepest descent in the objective function landscape. Higher derivatives (like the Hessian $\nabla^2_\theta J$) provide additional curvature information. In the following, we will for simplicity of notation abstract from the details of $J$. All we require is that our objective can be written in the form

$$J(\theta) = \sum_{j=0}^{N} J_j(y(t_j,\theta),\theta)$$

as a sum over the discrete time points $t_0 < \ldots < t_N$, where we w.l.o.g. hide dependence on data and experimental conditions. When we have multiple experimental conditions, experiments, or observables, one can simply sum over all of them inside the $J_j$. Then, we have for the derivative

$$\begin{aligned}
\frac{dJ}{d\theta}(\theta) &= \sum_{j=0}^{N} \left[ \frac{\partial J_j}{\partial \theta}(y(t_j,\theta),\theta) + \frac{\partial J_j}{\partial y}(y(t_j,\theta),\theta)\frac{dy}{d\theta}(t_j,\theta) \right] \\
&= \sum_{j=0}^{N} \left[ \frac{\partial J_j}{\partial \theta}(y(t_j,\theta),\theta) + \frac{\partial J_j}{\partial y}(y(t_j,\theta),\theta)\frac{\partial h}{\partial \theta}(x(t_j,\theta),\theta) + \frac{\partial J_j}{\partial y}(y(t_j,\theta),\theta)\frac{\partial h}{\partial x}(x(t_j,\theta),\theta)\frac{dx}{d\theta}(t_j,\theta) \right] \quad (3) \\
&= \sum_{j=0}^{N} \left[ \frac{\partial J_j}{\partial \theta} + \frac{\partial J_j}{\partial y}\frac{\partial h}{\partial \theta} + \frac{\partial J_j}{\partial y}\frac{\partial h}{\partial x}\frac{dx}{d\theta} \right],
\end{aligned}$$

where we used

$$\frac{dy}{d\theta} = \frac{\partial h}{\partial \theta} + \frac{\partial h}{\partial x}\frac{dx}{d\theta},$$

and dropping the obvious arguments at time points $t_j$. The interesting part here is the last summand which involves $\frac{dx}{d\theta}$, the derivative w.r.t. the parameter $\theta$ of the state $x$, which in turn is given only as the solution of an ODE. To compute $\nabla_\theta J$, there exist – apart from finite differences, which are often observed to be numerically unstable and computational expensive – basically two approaches, which we will shortly discuss.

#### 2.2 Forward sensitivity analysis

We can compute $\nabla_\theta J$ once we have the state sensitivities $s^x := \frac{dx}{d\theta}$, or equivalently the observable sensitivities $s^y := \frac{dy}{d\theta} = \frac{\partial h}{\partial x}s^x + \frac{\partial h}{\partial \theta}$. They can be obtained by differentiating the ODE (1) w.r.t. $\theta$, giving the forward sensitivity equations

$$\dot{s}^x(t,\theta) = \frac{\partial f}{\partial x}(x(t,\theta),\theta)s^x(x(t,\theta),\theta) + \frac{\partial f}{\partial \theta}(x(t,\theta),\theta), \quad s^x(0,\theta) = \frac{dx_0}{d\theta}(\theta).$$

These ODEs can be solved along with the original ODE, giving an enlarged system of ODEs

$$\begin{aligned}
&\dot{x} = f, && x(0) = x_0, \\
&\dot{s}_i^x = \tfrac{\partial f}{\partial x}s_i^x + \tfrac{\partial f}{\partial \theta_i}, && s_i^x(0) = \tfrac{dx_0}{d\theta_i}, \qquad i = 1,\ldots,n_\theta
\end{aligned}$$

of dimension $n_x(1+n_\theta)$, the solution of which is the state $x$ along with its sensitivities $s^x$.

#### 2.3 Adjoint sensitivity analysis

An alternative way of getting rid of the problematic term in (3) is as follows: We introduce an adjoint state $p : [t_0, t_N] \to \mathbb{R}^{n_x}$ playing the role of a Lagrange multiplier. Then, with $\frac{d\dot{x}}{d\theta} = \frac{\partial f}{\partial \theta} + \frac{\partial f}{\partial x}\frac{dx}{d\theta}$ according to the differential equation, we can write, by adding a zero-term,

$$\begin{aligned}
\frac{\partial J_j}{\partial y}\frac{\partial h}{\partial x}\frac{dx}{d\theta} &= \frac{\partial J_j}{\partial y}\frac{\partial h}{\partial x}\frac{dx}{d\theta} + \int_{t_j}^{t_{j+1}} p^T\left(\frac{d\dot{x}}{d\theta} - \frac{\partial f}{\partial \theta} - \frac{\partial f}{\partial x}\frac{dx}{d\theta}\right)dt \\
&\overset{IP}{=} \frac{\partial J_j}{\partial y}\frac{\partial h}{\partial x}\frac{dx}{d\theta} + \left(\lim_{t \nearrow t_{j+1}} p^T\frac{dx}{d\theta} - \lim_{t \searrow t_j} p^T\frac{dx}{d\theta}\right) - \int_{t_j}^{t_{j+1}} \left(\dot{p}^T + p^T\frac{\partial f}{\partial x}\right)\frac{dx}{d\theta}dt - \int_{t_j}^{t_{j+1}} p^T\frac{\partial f}{\partial \theta}dt,
\end{aligned}$$

$j = 0,\ldots,N-1$. Here, it becomes apparent that it is convenient to choose $p$ s.t. it satisfies $p(t) = 0$ for $t > t_N$, and on $(t_j, t_{j+1}]$ $p$ satisfies the backward differential equation

$$\dot{p} = -\frac{\partial f}{\partial x}^T p \quad \text{with initial condition} \quad p(t_{j+1},\theta) = \lim_{t \searrow t_{j+1}} p(t) - \frac{\partial J_{j+1}}{\partial y}\frac{\partial h}{\partial x}. \quad (4)$$

Then, in the above expression the first integral vanishes, and due to the re-initialization of the backward equation at time point $t_j$, the expression simplifies to

$$\begin{aligned}\frac{\partial J_j}{\partial y}\frac{\partial h}{\partial x}\frac{dx}{d\theta} &= \frac{\partial J_j}{\partial y}\frac{\partial h}{\partial x}\frac{dx}{d\theta} + p(t_{j+1},\theta)^T\frac{dx}{d\theta}(t_{j+1},\theta) \\ &\quad - (p(t_j,\theta)^T + \frac{\partial J_j}{\partial y}(y(t_j,\theta),\theta)\frac{\partial h}{\partial x}(t_j,x(t_j,\theta),\theta))\frac{dx}{d\theta}(t_j,\theta) - \int_{t_j}^{t_{j+1}} p^T\frac{\partial f}{\partial \theta}dt \\ &= p(t_{j+1},\theta)^T\frac{dx}{d\theta}(t_{j+1},\theta) - p(t_j,\theta)^T\frac{dx}{d\theta}(t_j,\theta) - \int_{t_j}^{t_{j+1}} p^T\frac{\partial f}{\partial \theta}dt.\end{aligned}$$

Summing over $j = 0,\dots,N-1$, simplifying the telescope sum, and adding $p(t_N)\frac{dx}{d\theta} = -\frac{\partial J_N}{\partial y}\frac{\partial h}{\partial x}\frac{dx}{d\theta}$, we get

$$\frac{dJ}{d\theta} = \sum_{j=0}^{N}\left[\frac{\partial J_j}{\partial \theta} + \frac{\partial J_j}{\partial y}\frac{\partial h}{\partial \theta}\right] - p(t_0,\theta)^T\frac{dx_0}{d\theta}(\theta) - \int_{t_0}^{t_N} p^T\frac{\partial f}{\partial \theta}dt. \tag{5}$$

This is independent of the state sensitivities $\frac{dx}{d\theta}$, except from those at time 0, which can however usually be easily computed. In turn, it now depends on the adjoint state $p$, whose dimension is $n_x$ and does not increase with the dimension of $\theta$. Thus, compared to forward sensitivity analysis, the computational effort for one gradient evaluation has changed from $n_\theta + 1$ systems of size $n_x$ to 2 systems of size $n_x$. In addition, a numerical quadrature is required, the effort for which is often negligible. Computation times may however also be affected by e.g. stiffness of the quadrature as well as the need to store results of the forward equation at intermediate points.

##### 2.4 Combining hierarchical optimization with adjoint sensitivities

In this manuscript, we assume the particular unscaled observation function

$$\tilde{y}(t,\theta,u) = \tilde{h}(x(t,\theta,u),\theta,u)$$

s.t. the former full (scaled) observation function becomes

$$y_{i_y}(t_{i_t},\theta,u_{i_u}) = h_{i_y}(x(t_{i_t},\theta,u_{i_u}) = s_{i_t,i_y,i_u}\cdot\tilde{h}_{i_y}(x(t_{i_t},\theta,u_{i_u}),\theta,u_{i_u}) + b_{i_t,i_y,i_u} \tag{6}$$

with scaling parameters $s$ and offset parameters $b$, to put the simulations on the same scale as the measurements. Then, simulations and data are related via

$$\bar{y}_{i_t,i_y,i_u} = y_{i_t,i_y,i_u} + \epsilon_{i_t,i_y,i_u},$$

$\epsilon$ encoding the (additive) noise model, which we here assume to be Gaussian independent additive (see main manuscript). Together, our negative log-likelihood objective function (2) thus reads

$$J(\theta,s,b,\sigma) = \sum_{i_t,i_y,i_u}\left[\log(2\pi\sigma^2_{i_t,i_y,i_u}) + \frac{(\bar{y}_{i_t,i_y,i_u} - (s_{i_t,i_y,i_u}\cdot\tilde{h}_{i_y}(t_{i_t},\theta,u_{i_u}) + b_{i_t,i_y,i_u}))^2}{\sigma^2_{i_t,i_y,i_u}}\right]. \tag{7}$$

For ease of notation, we summarize the static parameters as

$$w := (s,b,\sigma)$$

such that our overall parameter vector becomes $(\theta, w)$ with dynamic parameters $\theta$ and static parameters $w$. Further, we abstract from any condition dependence. Then, we can write

$$y(t,\theta,w) = h(\tilde{h}(x(t,\theta),\theta),w),$$

s.t. only the dynamic parameters are involved in the ODE simulation, where

$$\tilde{y}(t,\theta) = \tilde{h}(x(t,\theta),\theta)$$

denotes the mapping from the states $x$ to the unscaled observables $\tilde{y}$, which have to be scaled to $y$ using $w$ in order to allow for the comparison to the observed data. As in the discussion for the derivation of forward and adjoint sensitivity analysis above (there yet without static parameters), we write our objective function (7), highlighting only time dependence (with $N := n_t - 1$), as

$$J(\theta, w) = \sum_{j=0}^{N} J_j(\tilde{y}(t_j, \theta), \theta, w).$$

Note that here we added some extra $\theta$ parameters in $\tilde{h}$ and $J_j$ compared to (6), in case one does not treat all parameters not directly involved in the ODE simulation of $x$ as static parameters, but rather as dynamic parameters. This notation thus allows for more flexibility.

When we want to estimate the parameters, the most straightforward way is to treat both kinds of parameters equally, i.e. to optimize over the full vector $(\theta, w)$ simultaneously. This we refer to as the standard optimization approach. However, it can be favorable to use hierarchical optimization as introduced in the main manuscript on

$$\hat{J}(\theta) := J(\theta, w(\theta)), \quad \text{where } w(\theta) := \arg\min_{w} J(\theta, w).$$

Now, assuming $\hat{J}$ is differentiable, the gradient is given as

$$\begin{aligned}\frac{d\hat{J}}{d\theta} &= \frac{\partial J}{\partial \theta} + \frac{\partial J}{\partial w}\frac{dw}{d\theta} \\ &= \sum_{j=0}^{N}\left[\frac{\partial J_j}{\partial \theta} + \frac{\partial J_j}{\partial \tilde{y}}\frac{d\tilde{y}}{d\theta} + \frac{\partial J_j}{\partial w}\frac{dw}{d\theta}\right] \\ &= \sum_{j=0}^{N}\left[\frac{\partial J_j}{\partial \theta} + \frac{\partial J_j}{\partial \tilde{y}}\frac{\partial \tilde{h}}{\partial \theta} + \frac{\partial J_j}{\partial \tilde{y}}\frac{\partial \tilde{h}}{\partial x}\frac{dx}{d\theta} + \frac{\partial J_j}{\partial w}\frac{dw}{d\theta}\right].\end{aligned} \tag{8}$$

Here, assuming that the inner optimization problem was solved s.t.

$$\frac{\partial J}{\partial w} = 0 \tag{9}$$

(necessary for a local minimum in the interior of the domain of definition, and for our applications always the case), the last term vanishes and we do not need to compute the terms $\frac{dw}{d\theta}$, as our gradient reduces to

$$\frac{d\hat{J}}{d\theta} = \sum_{j=0}^{N}\left[\frac{\partial J_j}{\partial \theta} + \frac{\partial J_j}{\partial \tilde{y}}\frac{\partial \tilde{h}}{\partial \theta} + \frac{\partial J_j}{\partial \tilde{y}}\frac{\partial \tilde{h}}{\partial x}\frac{dx}{d\theta}\right]. \tag{10}$$

Then, in one optimizer iteration, for given $\theta$, we can first simulate the observables $y(\theta)$ for the required time points and experimental conditions, then compute the optimal $w(\theta)$, and then compute the objective $\hat{J}$ as well as potentially derivatives to guide optimization. This was done using forward sensitivity analysis in Loos et al. [2018]. While we will not go into details here, the basic algorithm works as follows:

1. $\tilde{y}, d_\theta \tilde{y}$ = simulate forward given $\theta$,
2. $w, d_\theta w$ = compute optimal static parameters given $\tilde{y}, \mathcal{D}$,
3. $\hat{J}, \nabla \hat{J}$ = compute likelihood given $w, d_\theta w, \mathcal{D}, \tilde{y}, d_\theta \tilde{y}$.

Here, in the first step the unscaled states and their sensitivities w.r.t. $\theta$ are simulated, and in the third step all derivatives are gathered according to (10) to give the full gradient. Here, an ODE simulation is involved only in the first step.

However, the question remained open whether the hierarchical approach can also be combined with adjoint sensitivities, which is of great interest for large-scale models. The main problem here is that, as we see in equation (4), in general the adjoint state depends on the data and the full simulated observables $y$ via the residuals $J_j$. But unlike $\tilde{y}$, $y$ depends also on the static parameters $w$. So, we cannot first simulate the ODE and then compute optimal static parameters and the objective along with its gradient, as done in the forward approach. Nonetheless, combining the hierarchical approach with adjoints turns out to be not that difficult: Again, there is only one annoying term in (10) involving the state sensitivities $\frac{dx}{d\theta}$, which depends on an ODE, all the other terms are in practice readily computed. In this term, it does not matter how $w$ was computed, so it can be treated as constant and $h$ as a function of only $x$ and $\theta$. So, replacing $\frac{\partial h}{\partial x}$ by $\frac{\partial \tilde{h}}{\partial x}$, we see that we can again apply the adjoint equation (4) to get an adjoint formulation comparable to (5), here taking the form

$$\frac{d\hat{J}}{d\theta} = \sum_{j=0}^{N} \left[ \frac{\partial J_j}{\partial \theta} + \frac{\partial J_j}{\partial \tilde{y}} \frac{\partial \tilde{h}}{\partial \theta} \right] - p(t_0, \theta)^T \frac{dx_0}{d\theta}(\theta) - \int_{t_0}^{t_N} p^T \frac{\partial f}{\partial \theta} dt.$$

Then, in practice we can proceed as follows: First, we simulate the unscaled observables $\tilde{y}$, then compute optimal static parameters $w$, and then use both to compute the likelihood $\hat{J}$ as well as to simulate the adjoint state $p$ which allows us to compute $\nabla \hat{J}$. In algorithmic form:

1. $\tilde{y}$ = simulate forward given $\theta$,
2. $w, d_\theta w$ = compute optimal static parameters given $\tilde{y}, \mathcal{D}$,
3. compute $\hat{J}$ given $\tilde{y}, w, \mathcal{D}$,
4. simulate adjoint state $p$ given $\tilde{y}, w, d_\theta w, \mathcal{D}$ to get $\nabla \hat{J}$.

This algorithm requires no additional ODE simulations compared to the standard approach. Moreover, since the number of parameters is reduced, we can hope for reduced computation times, and in particular that the problem is easier to optimize.

It should be noted that since the inner subproblem was assumed to be solved exactly in this chapter in the sense of (9), s.t. the gradient reduces to

$$\nabla \hat{J}(\theta) = \frac{\partial J}{\partial \theta}(\theta, w(\theta)),$$

all expressions involving $\frac{\partial J}{\partial w}$ vanish from the gradient, so that (8) effectively reduces to (3) with the additional application of rescalings to $\tilde{h}$ in $J_j$. Also, then $\frac{dw}{d\theta}$ does not need to be evaluated. From a practical perspective, this implies that, once computed, we can treat the static parameters $w$ as constant when computing $\frac{d\hat{J}}{d\theta}$ using the adjoint approach. Thus, we can easily re-use existing software implementations that allow to pass parameters $\theta$ the gradient shall be computed along, as well as constants (primarily used to express differential experimental conditions) along which no gradients are supposed to be evaluated.

If the inner subproblem is not solved exactly in the sense of (9), adding these terms to the gradient $\frac{d\hat{J}}{d\theta}$ involves some additional effort and cannot be addressed by the proposed approach.

#### 2.5 Properties of the different optimization approaches

In Table 1, the properties of the diverse optimization approaches discussed above, standard vs. hierarchical and forward vs. adjoint sensitivities, are listed. *Scalable* means that the approach scales well with the number of parameters. *Handle static parameters efficiently* means that the handling of additional static parameters is possible without considerable additional computational overhead. *FIM uncertainty assessment* means that the Fisher Information Matrix can be computed in order to assess parameter uncertainty. *Least-squares based optimization* means that (non-linear) least-squares optimizers can be used, which are based on the observable residuals, in contrast to *objective based optimization*, which only requires the full objective function. Finally, *improved convergence* means that the approach has been observed to converge more often to the global optimum than the other approaches.

Table 1: Overview of optimization approaches

| | scalable | handle static parameters efficiently | FIM uncertainty assessment | objective based optimization | least-squares based optimization | improved convergence |
|---|---|---|---|---|---|---|
| standard, forward | | | ✓ | ✓ | ✓ | |
| standard, adjoint | ✓ | | | ✓ | | |
| hierarchical, forward | | ✓ | ✓ | ✓ | ✓ | ✓ |
| hierarchical, adjoint | ✓ | ✓ | | ✓ | | ✓ |

### 3 Analytic formulas for scaling, offset, and noise parameters for additive Gaussian measurement noise

In this section, we derive general formulas for the scaling, offset, and noise parameters assuming additive Gaussian noise. We consider the observation function (6) and objective function (7) from the previous chapter.

In general, scaling parameters $s$, offset parameters $b$, and noise parameter $\sigma$ can be shared among various data points, and either need to be estimated or sometimes can be fixed externally. Sharing parameters translates to using e.g. the same rescaling for all data points in a time series, or generated under the same conditions. To define which time points, observables, and experimental conditions share a scaling, offset, or noise parameter, one can define

$$I^s_\alpha, I^b_\beta, I^\sigma_\gamma \subset \mathbb{N}^{n_t}_+ \times \mathbb{N}^{n_y}_+ \times \mathbb{N}^{n_u}_+$$

for $\alpha = 1, \ldots, n_s$, $\beta = 1, \ldots, n_b$, $\gamma = 1, \ldots, n_\sigma$, where $n_s, n_b, n_\sigma$ denote the numbers of scaling, offset, and noise parameters respectively, s.t. each data point belongs to exactly one $I^s_\alpha, I^b_\beta, I^\sigma_\gamma$. Then, all parameters $s_{i_t,i_y,i_u}$ for which the indices $i = (i_t, i_y, i_u)$ are contained in the same group $I^{i_s}_s$ share the same scaling parameters, similarly for offset and noise parameters.

A common assumption in the following will be that first,

$$\{I^s_\alpha\}_\alpha = \{I^b_\beta\}_\beta ,$$

i.e. the sets $I^s = I^b$ coincide, and that second

$$\forall \alpha \exists \gamma : I^s_\alpha \subset I^\sigma_\gamma,$$

i.e. data points sharing the scale parameter (and thus also the offset parameter) share also the noise parameter. This we will also refer to as the resolution of $\sigma$ being coarser than that of $s$ and $b$.

These two assumptions will be required only when all parameters are to be optimized. We will call variables fixed, if they are not to be optimized, but specified to some fixed value externally, or optimized as a dynamic parameter. This will be made explicit in the respective subsections.

For ease of notation, we write in the following

$$\bar{y}_i = s_i \cdot \tilde{h}_i + b_i + \epsilon_i,$$

where $i \in I$ is an arbitrary index over e.g. time points, observables, replicates, and experimental conditions, and $\tilde{h}_i = \tilde{h}_i(t_i, \theta, u_i)$ denotes the unscaled observable. So, in the following, we consider the negative log-likelihood

$$J(\theta, b, c, \sigma^2) = \frac{1}{2} \sum_{i \in I} \left[ \log(2\pi\sigma_i^2) + \frac{(\bar{y}_i - (s_i \cdot \tilde{h}_i + b_i))^2}{\sigma_i^2} \right]$$

for minimization, assuming a Gauss noise variable, see (2). We can decompose the objective function $J$ to sums over indices of interest, which we will w.l.o.g. do in the following, i.e. in the following sections, we only consider the part of the sum relevant for a given parameter.

##### 3.1 Scalings and offsets

We assume that the resolution of the scalings $s$ and offsets $b$ coincide, and that either $\sigma^2$ is not to be optimized, or its distribution is coarser than that of the scalings. So, we consider a reduced sum

$$J(\theta, s, b, \sigma^2) := \frac{1}{2}\sum_i \left[\log(2\pi\sigma_i^2) + \frac{(\bar{y}_i - (s\cdot\tilde{h}_i + b))^2}{\sigma_i^2}\right],$$

where we assume that either $\sigma^2$ is not to be optimized as a static parameter, or, if they are to be optimized as static parameters. In that case imagine $\sigma_i^2 = 1$ for all $i$, as it will then drop out of the formulas here and can be determined afterwards.

To find optimal parameters, we have the necessary conditions that the derivatives w.r.t. said parameters disappear:

$$0 = \partial_s J = -\sum_i \frac{\bar{y}_i - (s\tilde{h}_i + b)}{\sigma_i^2}\tilde{h}_i \Rightarrow s = \left(\sum_i \frac{\tilde{h}_i^2}{\sigma_i^2}\right)^{-1}\left(\sum_i \frac{(\bar{y}_i - b)\tilde{h}_i}{\sigma_i^2}\right)$$

and

$$0 = \partial_b J = -\sum_i \frac{\bar{y}_i - (s\tilde{h}_i + b)}{\sigma_i^2} \Rightarrow b = \left(\sum_i \frac{1}{\sigma_i^2}\right)^{-1}\left(\sum_i \frac{\bar{y}_i - s\tilde{h}_i}{\sigma_i^2}\right).$$

If either scaling or offset is fixed, we are done by inserting that value (in particular 1 or 0 assuming absolute data) in the respective formula for the other one. Note that also different values for the fixed variables for different data points would be possible. If both are to be optimized, we proceed by inserting the first equation into the second one, giving

$$\left[1 - \left(\sum_i \frac{1}{\sigma_i^2}\right)^{-1}\left(\frac{\sum_i \frac{\tilde{h}_i}{\sigma_i^2}\sum_j \frac{\tilde{h}_j}{\sigma_j^2}}{\sum_i \frac{\tilde{h}_i^2}{\sigma_i^2}}\right)\right] b = \left(\sum_i \frac{1}{\sigma_i^2}\right)^{-1}\left(\sum_i \frac{\bar{y}_i}{\sigma_i^2} - \frac{\sum_i \frac{\bar{y}_i\tilde{h}_i}{\sigma_i^2}\sum_j \frac{\tilde{h}_j}{\sigma_j^2}}{\sum_i \frac{\tilde{h}_i^2}{\sigma_i^2}}\right).$$

If the factor on the left-hand side is non-zero, we find

$$b = \frac{\left(\sum_i \frac{1}{\sigma_i^2}\right)^{-1}\left(\sum_i \frac{\bar{y}_i}{\sigma_i^2} - \frac{\sum_i \frac{\bar{y}_i\tilde{h}_i}{\sigma_i^2}\sum_j \frac{\tilde{h}_j}{\sigma_j^2}}{\sum_i \frac{\tilde{h}_i^2}{\sigma_i^2}}\right)}{1 - \left(\sum_i \frac{1}{\sigma_i^2}\right)^{-1}\left(\frac{\sum_i \frac{\tilde{h}_i}{\sigma_i^2}\sum_j \frac{\tilde{h}_j}{\sigma_j^2}}{\sum_i \frac{\tilde{h}_i^2}{\sigma_i^2}}\right)}. \tag{11}$$

Otherwise, our simulations are not diverse enough to allow the identification of both $s$ and $b$, in which case we can simply set $b = 0$. However, if such a case is detected, a warning to the user is a good idea, as it indicates that revising the data and static parameters would be a good idea. In either case, we can then use $b$ to compute $s$ as in the previous formula.

Inspection of the second derivative shows that

$$\partial_s^2 J = \sum_i \frac{\tilde{h}_i^2}{\sigma_i^2} \geq 0, \quad \text{and} \quad \partial_b^2 J = \sum_i \frac{1}{\sigma_i^2} > 0,$$

so both are obviously (global) minima.

##### 3.2 Noise parameters

We consider the reduced objective function

$$J(\theta, s, b, \sigma^2) = \frac{1}{2}\sum_i \left[\log(2\pi\sigma^2) + \frac{(\bar{y}_i - (s_i\tilde{h}_i + b_i))^2}{\sigma^2}\right].$$

This sum will typically involve multiple sums of the type discussed for the scalings and offsets. Here, it does not matter where the values for $s_i$ and $b_i$ come from. We find

$$0 = \partial_{\sigma^2} J = \frac{1}{2}\sum_i \left[\frac{1}{\sigma^2} - \frac{(\bar{y}_i - (s_i\tilde{h}_i + b_i))^2}{(\sigma^2)^2}\right] \Rightarrow \sigma^2 = \left(\sum_i 1\right)^{-1}\left(\sum_i (\bar{y}_i - (s_i\tilde{h}_i + b_i))^2\right).$$

Inspection of the second derivative shows that

$$\begin{aligned}\partial^2_{\sigma^2} J &= \frac{1}{2}\sum_i \frac{2(\bar{y}_i - (s_i\tilde{h}_i + b_i))^2 - \sigma^2}{(\sigma^2)^3} \\ &= \frac{1}{2}\frac{2\sum_i(\bar{y}_i - (s_i\tilde{h}_i + b_i))^2 - \sum_i(\bar{y}_i - (s_i\tilde{h}_i + b_i))^2}{(\sigma^2)^3} \\ &= \frac{1}{2}\frac{\sum_i(\bar{y}_i - (s_i\tilde{h}_i + b_i))^2}{(\sigma^2)^3} = \frac{1}{2}\frac{\sum_i 1}{(\sigma^2)^2} > 0,\end{aligned}$$

confirming we have a minimum.

A problem arises when for the optimal noise parameter holds $\sigma(\theta) = 0$. This indicates that all (rescaled) simulation and data that are relevant for the given $\sigma$ are a perfect match (easily possible e.g. when there is just one data point per noise parameter). In this case, we would have a singularity for the objective function, which is undesirable as it does not allow to infer the other parameters. Such constellations should be filtered out beforehand. If this is not possible, a practical solution would be to limit $\sigma > \sigma_0$ for some small minimum $\sigma_0 > 0$. For the models and data used in this study, this was not an issue.

##### 3.3 Objective gradient used for optimization

The objective function gradient then used in optimization (assuming $s, b, \sigma \notin \theta$, otherwise those derivatives need to be added) is given specifically by

$$\nabla \hat{J}(\theta) = \partial_\theta J(\theta, s(\theta), b(\theta), \sigma(\theta)) = \sum_i \frac{\bar{y}_i - (s_i\tilde{h}_i + b_i)}{\sigma_i^2} s \frac{d\tilde{h}_i}{d\theta},$$

as one easily verifies. While this expression still contains the sensitivities of states w.r.t. $\theta$, we employed a reformulation using adjoint sensitivity analysis, as explained in the previous chapter, see also Stapor et al. [2018] for the explicit formulas in the Gaussian case.

### 4 Implementation and computational analysis

##### 4.1 Measurements for model calibration

We used the data file `MCLP v1.1` from the MCLP website (`https://tcpaportal.org/mclp/#/download`, downloaded on 23.03.2017). 96 cell-lines of the processed CCLE measurements from Froehlich et al. [2018] were taken.

##### 4.1.1 Mapping protein measurements to the ODE model

To map the measured proteins from, e.g. MCLP, we added an observable as the sum over all states that included this specific protein, weighted by their stoichiometric coefficients. Phosphorylation site-specific antibodies were assumed to bind any modeled state of the given protein carrying the respective phosphorylation (irrespective of any other modification). Antibodies not specific for any phospho-site were assumed to bind any modeled state of this protein, modified or unmodified.

#### 4.2 Model simulation

Model simulation and gradient computation was performed using AMICI (`https://github.com/ICB-DCM/AMICI/`, commit 974093). The SBML model was imported using the AMICI Matlab interface (Matlab, Mathworks, versions R2017a or R2017b) and simulated using either the Matlab interface (for fmincon optimization) or the C++ interface (other optimizers). For computing the gradient $\nabla J$ of the objective function, we employed the adjoint approach as implemented in AMICI.

All measurements were considered steady-state measurements and the model was simulated from $t_0 = 0, x_0 = 0$ until $t = 10^8$, which showed to be sufficient for equilibration in previous studies.

#### 4.3 Comparison of computation time for forward and adjoint sensitivities

We estimated the speedup using adjoint sensitivities by randomly sampling 100 combinations of parameters and experimental conditions. To reduce computation time, we assumed that the calculation of forward sensitivities scales linearly with the number of parameters and interpolated the evaluation time from the gradient calculation for a randomly sampled subset of 20 parameters. We observed an estimated mean speedup of $\sim 1.5 \times 10^3$. One full gradient evaluation using forward sensitivities would need an estimated computation time of $\sim 6 \times 10^3$ hours.

#### 4.4 Optimization objective and gradient computation

With all optimizers we used an objective of the form of (2). The model was simulated for each experimental condition and for the standard setting, the log-likelihood as well as, where required, its gradient as provided by AMICI were aggregated.

In the hierarchical optimization setting, simulations were performed the same way, but then optimal static parameters were computed, the observables rescaled and the log-likelihood was computed. For computing the gradient $\nabla J$ of the objective function, we employed the adjoint approach as explained above. It was, for ease of implementation, that the forward simulation was performed twice, once before computing the optimal static parameters, and afterwards again in order to compute the adjoint state. This was because intermediate results from the forward simulation are required for the backward simulation, which we did not store. So, we had a little computational overhead compared to the theoretically most efficient algorithm depicted in Figure 1 of the main manuscript.

#### 4.5 Parameter optimization

##### 4.5.1 Parameter optimization settings

For parameter optimization, we used log-transformed parameters except for the offsets, which may also take negative values. Initial parameters were sampled within the interval $[-2, 2]$ ($[-10^2, 10^2]$ for offsets) and for optimization, the parameters were bound by $[-5, 3]$ ($[-10^3, 10^3]$ for offsets). Wider bounds sometimes led to numerical problems, because of an increased number of parameters for which the objective function or the gradients were not evaluable.

We chose the parameter bounds to be wider than the sampling interval, to avoid problems that occur, especially with Ipopt, when the starting parameters are too close to the bounds.

Initially, we encountered premature stopping of the optimizers due to failure to evaluate the objective function at certain parameter values. Therefore, we implemented an automated integration tolerance relaxation scheme. In case of integration failures, we repeated the respective simulation with absolute and relative tolerances relaxed by a factor of 100, starting from an initial absolute tolerance of $10^{-16}$ and relative tolerance $10^{-8}$. This process was repeated for three times.

##### 4.5.2 Parameter optimization fmincon

For parameter optimization, objective function for hierarchical and standard optimization was implemented in Matlab (Supplemental data, script path. Parameter estimation was performed using PESTO (`https://github.com/ICB-DCM/PESTO/`, commit 1e30f9). Matlab code was compiled using the Matlab compiler toolbox and run on compute nodes equipped with either: AMD Opteron processors with nominal frequency 2.4 GHz, 64 cores and 512 GB / AMD Opteron processors with nominal frequency 2.4 GHz, 64 cores and 256 GB / AMD Opteron processors with nominal frequency 1.7 GHz, 48 cores and 192 GB. Model simulations were parallelized across 32 cores using Matlab `parfor`. Individual local optimizations were run independently. Fmincon was supplied with gradients as described above and called with the following options: *MaxIter*: 100, *TolX*: $10^{-8}$, *TolFun*: 0, *MaxFunEvals*: $10^7$, *algorithm*: `interior-point`, *GradObj*: `on`. We used the same AMICI version and settings as for the optimizers described in the following section. However, due to different hardware, the reported run times for fmincon cannot be compared directly to those of the other optimizers.

##### 4.5.3 Parameter optimization Ipopt, SUMSL, Ceres

For parameter optimization, we used the following optimizers and settings:

- Ipopt (Wächter and Biegler [2006], `https://www.coin-or.org/Ipopt/`, version 3.12.9 with COIN-HSL version 2015.06.23 [HSL, 2019])
  Default settings were used except for: *max_iter*: 150, *hessian_approximation*: `limited-memory`, *limited_memory_update_type*: `bfgs`, *tol*: $10^{-9}$, *acceptable_iter*: 1, *acceptable_tol*: $10^{20}$, *acceptable_obj_change_tol*: $10^{-12}$, *watchdog_shortened_iter_trigger*: 0.
- Ceres (Agarwal et al. [2019], `http://ceres-solver.org/`, version 1.13.0)
  We used the "General Unconstrained Minimization" algorithm with default settings except for: *max_num_iterations*: 150.
- SUMSL (Gay [1983], `http://netlib.org/toms/`, file `611.gz`)
  SUMSL is originally provided as Fortran code. We used `f2c` (Fortran to C Translator, version 20100827) to convert this to C++ code. To allow for parallel usage of SUMSL, all `static` variables were changed to `static __thread`. Default settings were used except for: Maximum number of objective function evaluations (*mxfcal*): $10^8$.

Ceres and SUMSL do not natively support parameter bounds. Therefore, we used a naive implementation, reporting failure to evaluate objective function to the optimizer in case the proposed parameters where outside the specified bounds.

We interfaced above-mentioned optimizers using a custom C++ parameter estimation library parPE (`https://github.com/ICB-DCM/parPE/`, commit `0e0fb1`) which provides a generalized objective function for AMICI models as described above and allows for its parallel evaluation. parPE was linked with the AMICI-generated model C++ codes to generate executables for parameter estimation and model simulation.

Parallelization at the level of individual simulations corresponding to individual experimental conditions was realized using the IBM message passing interface (MPI) library in a master-worker architecture. The respective optimizer was run on the master process and for objective function evaluation, batches of simulation jobs were sent to workers for evaluation. Aggregation of individual likelihoods and gradients, as well as where applicable, the computation of optimal static parameters were performed on master process. Different local optimizations for a given setting were either run independently or in parallel. In the case of parallel optimizer runs, MPI-based parallelization of simulations was combined with pthreads-parallel optimizer runs.

This code was run on the SuperMUC phase2 supercomputer at the Leibniz Supercomputing Centre (Leibniz-Rechenzentrum, LRZ) of the Bavarian Academy of Sciences and Humanities, Garching, Germany. The employed computes nodes were equipped with Haswell Xeon Processor E5-2697 v3 processors with nominal frequency 2.6 GHz, 28 cores per node and 64GB RAM per node. Compute jobs were submitted using IBM LoadLeveler using either 128 nodes (3576 cores) for parallel runs of 20 local optimizations within one job, or as 20 individual jobs using 10–12 nodes (280–336 cores).

#### 4.6 Data and code availability

All simulation, parameter estimation, data analysis and visualization codes are provided as Supplementary Code at http://doi.org/10.5281/zenodo.2592186. This archive additionally includes a human- and machine-readable definition of the parameter estimation problems considered in this manuscript in the PEtab format (https://github.com/ICB-DCM/PEtab). Intermediary data and result files are available at http://doi.org/10.5281/zenodo.2593839.

#### 4.7 Simulation study

##### 4.7.1 Synthetic data generation

We used synthetic absolute and relative viability data to evaluate the loss of information using relative data and the differences in optimizer performance. The data was simulated using parameters from a former optimization run, where viability and protein measurements were used. Noise was added to the simulated data according to the estimated sigma parameter for the viability data:

Notation:

$\hat{\cdot}$ - estimates from optimization with viability + protein data

$\hat{\theta}$ - dynamic parameters

$\hat{s}$ - scalings

$\hat{\sigma}_{\text{viability}}$ - sigma for viability data

With this notation, the synthetic and noisy relative data was generated by:

$$\bar{y}_{\text{rel}} = h(\hat{\theta}, \hat{s}) + \epsilon_{\text{rel}}, \quad \epsilon_{\text{rel}} \sim \mathcal{N}(0, \hat{\sigma}^2_{\text{viability}})$$

To get synthetic absolute data, the simulated relative data was first divided by the estimated scalings $\hat{s}$ to make them absolute and then noise was added:

$$\bar{y}_{\text{abs}} = \frac{h(\hat{\theta}, \hat{s})}{\hat{s}} + \epsilon_{\text{abs}}, \quad \epsilon_{\text{abs}} \sim \mathcal{N}\left(0, \left(\frac{\hat{\sigma}_{\text{viability}}}{\hat{s}}\right)^2\right)$$

##### 4.7.2 Parameter estimation

Parameters were estimated for three different settings:

1. Using $\bar{y}_{\text{abs}}$ as synthetic data
2. Using $\bar{y}_{\text{rel}}$ as synthetic data with standard optimization
3. Using $\bar{y}_{\text{rel}}$ as synthetic data with hierarchical optimization

Each time, Ipopt was used with 20 initial parameters and 150 maximal iterations. Each setting used the same initial dynamic parameters.

###### 4.7.3 Making relative data absolute

The relative simulations were made absolute by:

$$\tilde{h} = \frac{y_{\text{rel, sim}}(\theta^*, s^*)}{s^*},$$

where $^*$ indicates the here optimized parameters. As measurements, the absolute measurements $\bar{y}_{\text{abs}}$ were used. So the correlations were computed by:

$$corr(\bar{y}_{\text{abs}}, \tilde{h})$$

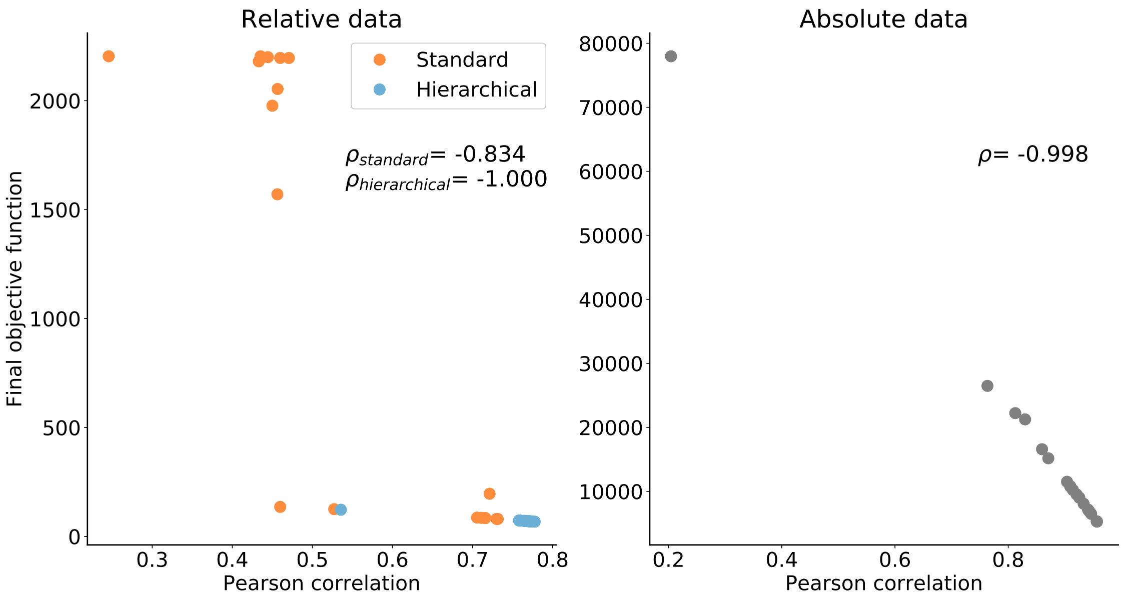

Figure S1: Scatter plot for final objective function values and correlation between model simulation and measurement for optimization with relative (left) and absolute data (right). Optimization was performed using simulated data. Related to main Figure 2.

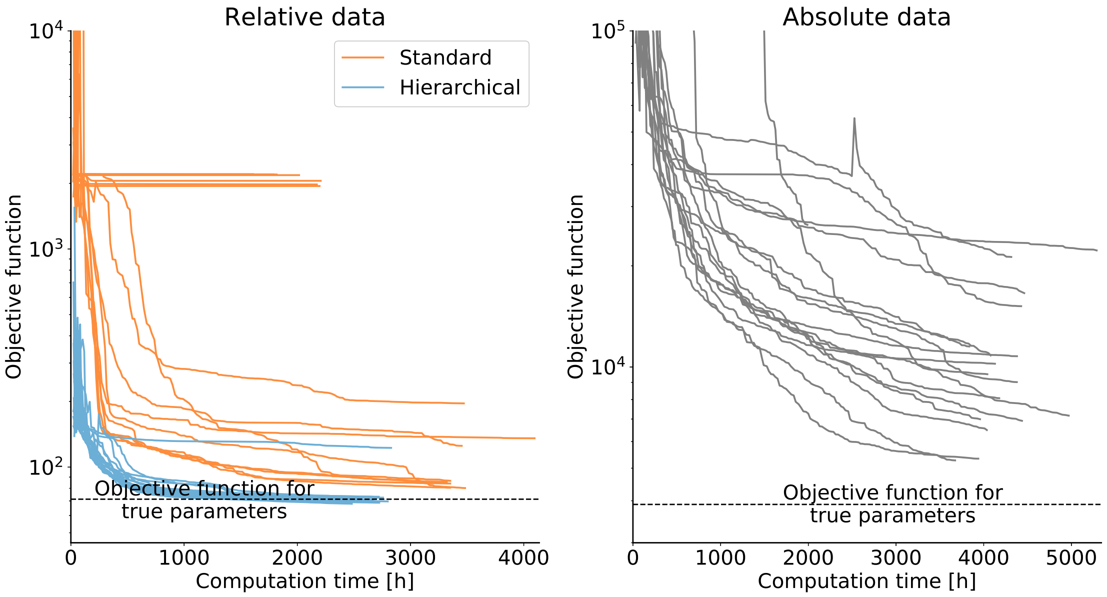

Figure S2: Optimizer trajectories for the different optimization settings. Dashed line shows objective function reached with true parameters. Optimization was performed using simulated data. Related to main Figure 2.

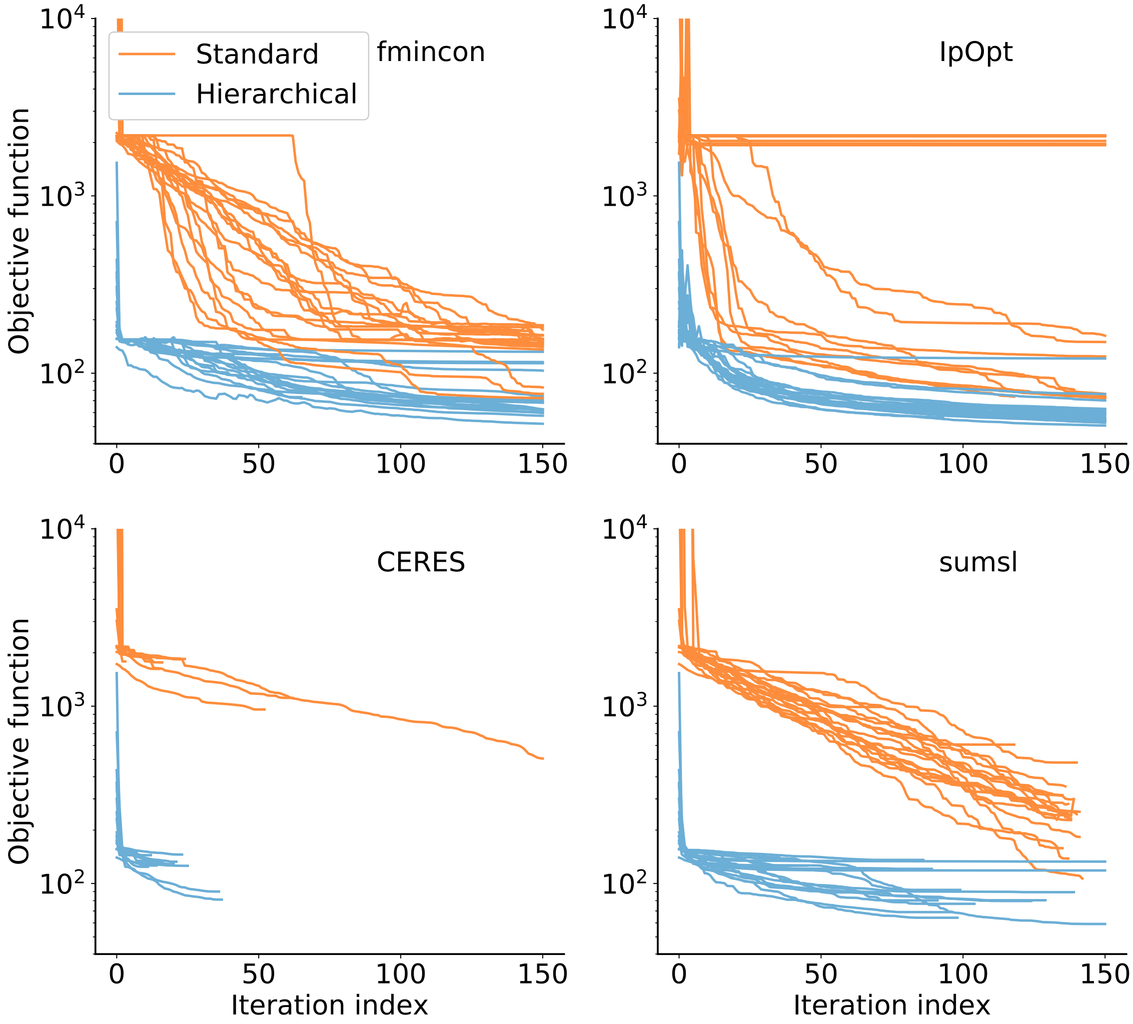

Figure S3: Optimizer trajectories for standard and hierarchical approach using *dataset 1* with Ipopt, fmincon, Ceres and sumsl. Optimization was performed using *dataset 1*. Related to main manuscript Figure 3.

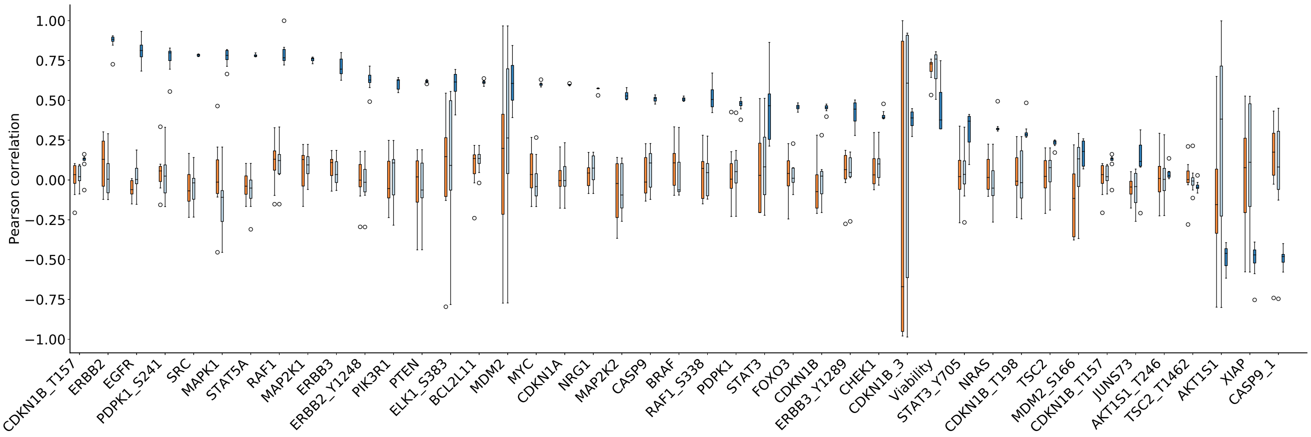

Figure S4: Pearson correlation for the best local optimization using standard and hierarchical optimization with *dataset 1&2*. All observables with more than 2 data points are shown. Related to main manuscript Figure 4.
